# Multi-omics analyses of drug repurposing reveal Acebutolol and Amiloride for osteoporosis treatment

**DOI:** 10.1101/2022.05.03.490380

**Authors:** Dan-Yang Liu, Jonathan Greenbaum, Qiao-Rong Yi, Shuang Liang, Yue Zhang, Jia-Chen Liu, Xiang-He Meng, Hong-Mei Xiao, Yun Deng, Li-Jun Tan, Hong-Wen Deng

## Abstract

Osteoporosis is a metabolic bone disease that occurs during aging, characterized by low bone mineral density (BMD) and a high risk of trauma fracture. While current pharmacological interventions provide symptomatic benefits, they are unsatisfactory and have major side effects. In this study, we used multi-omics data and drug similarity to construct osteoporosis driver signaling networks (ODSN) and drug functional networks (DFN), respectively. By integrating ODSN and DFN with treatment transcriptional responses, we observed 8 drugs that demonstrated strong targeting effects on ODSN. Mendelian Randomization analysis determines the causal effect on BMD using cis-eQTLs of the drug targets and BMD GWAS data. The findings suggested Acebutolol and Amiloride may increase BMD, while Acenocoumarol, Aminocaproic acid and Armodafinil may enhance bone loss. Zebrafish experiments experimentally showed Acebutolol hydrochloride and Amiloride hydrochloride had significant protective effects on osteoporosis in zebrafish embryos induced by Dexamethasone. Also, Acenocoumarol reduced bone mineralization compared with the control group. The findings suggest that the hypertension drugs Acebutolol and Amiloride warrant further investigation in functional mechanistic experiments to evaluate their effectiveness for osteoporosis treatments.

## 1. Introduction

Osteoporosis is a worldwide public health problem characterized by decreased bone mass and deteriorated microarchitectural, increased fragility fracture tendency ^1^. The prevalence of osteoporosis and fractures will increase rapidly as the global population continues to age ^2^. It was estimated that among the 27 countries of the European Union, 25 million women and 5.5 million men suffered from osteoporosis in 2010, and the economic burden of the incident and prior fragility fractures was estimated at EUR 37 billion ^3^. In China, the standardized prevalence of osteoporosis is expected to increase from 5.04% to 7.46% in males aged ≥ 50 years and from 26.28% to 39.19% in females aged ≥ 50 years over the duration of time from 1990 to 2050 ^4^. Hence, osteoporosis-related fractures result in a huge economic and societal burden in terms of time and expenditure.

Current pharmacological interventions for osteoporosis are categorized as either antiresorptive, which decreases the rate of bone resorption by osteoclasts, or anabolic, which stimulates bone formation by osteoblasts. Antiresorptive drugs include bisphosphonates, RANK ligand inhibitors (Denosumab) ^6^, and selective estrogen receptor modulators. Anabolic treatments are primarily derived from parathyroid hormone (PTH) analogues (Teriparatide and Abaloparatide) ^7^. Despite the remarkable advancements in our understanding of disease pathogenesis, these medications have several limitations ^8,9^. For example, bisphosphonates have been associated with atypical femur fractures, denosumab has been linked with musculoskeletal pain and hypercholesterolemia, and the duration of treatment with PTH analogues is limited due to the risk of osteosarcoma ^10^. Thus, identifying clinically meaningful novel treatments is still necessary for individuals at high risk of osteoporosis.

Given that novel traditional drug development is an extremely expensive and time-consuming process, repurposing drugs that were previously approved for other clinical outcomes is an attractive technique to potentially reduce costs and shorten the development timeline ^11^. Various computational strategies have been explored for drug repurposing such as literature mining, molecular docking, genome-wide association, pathway and signature mapping, and retrospective clinical analysis using electronic health records ^12^. Network-based approaches have proven especially useful to elucidate essential biochemical interact molecules in biological systems, improve the performance of the network-based algorithms and signaling system dynamics, and identify potential drug targets ^13,14^. Recently, Huang *et al* proposed a dysregulated driver signaling network identification and drug functional network (DSNI-DFN) pipeline, which is an intuitive system biology-based computational approach that integrates multi-omics profiles from the same samples to identify signaling networks that drive disease, and further evaluates the targeting effects of existing drugs on those networks ^15^. The integration of multi-omics data may detect molecular and genomic factors/mechanisms underlying the pathogenesis of disease in a powerfully and comprehensive understanding, which may represent the most critical therapeutic targets ^16^. In its original implementation, DSNI-DFN was successfully used to repurpose drugs of the cardiac glycoside family (e.g., digoxin) for treatment of medulloblastoma ^15^.

In the present study, we applied the DSNI-DFN approach to integrate multi-omics profiles from the same individuals (genome, transcriptome, methylome, metabolome) with interactome and pharmacogenomics information for the prediction of potential drug repurposing candidates which may be effective in the prevention/treatment of osteoporosis. Mendelian Randomization (MR) analyses were then used to evaluate the causal relationship between expression of the target genes for drug candidates and bone mineral density (BMD). Finally, a zebrafish model was implemented to validate the effects of the identified drugs on bone mass. This research innovatively applied the DSNI-DFN method and provides novel potential drug repurposing candidates for osteoporosis treatment.

## 2. Materials and methods

We screened novel drug candidates for osteoporosis by DSNI-DFN and Mendelian Randomization analysis based on the following steps (Figure 1): First, we identified driver signaling networks from a multi-omics dataset specifically constructed for osteoporosis research. Second, we used a similarity fusion approach to integrate drug similarity information from chemical structures, *in vitro* experiments, text-mining, and gene expression, and further obtained drug functional modules. Third, we ranked all the drugs by predicting the targeting effects on the osteoporosis driver signaling network. Then, we assessed the potential causal association between drug targets and BMD by MR analysis. Finally, we empirically tested and verified the drug effect using a zebrafish model.

**Figure 1.**
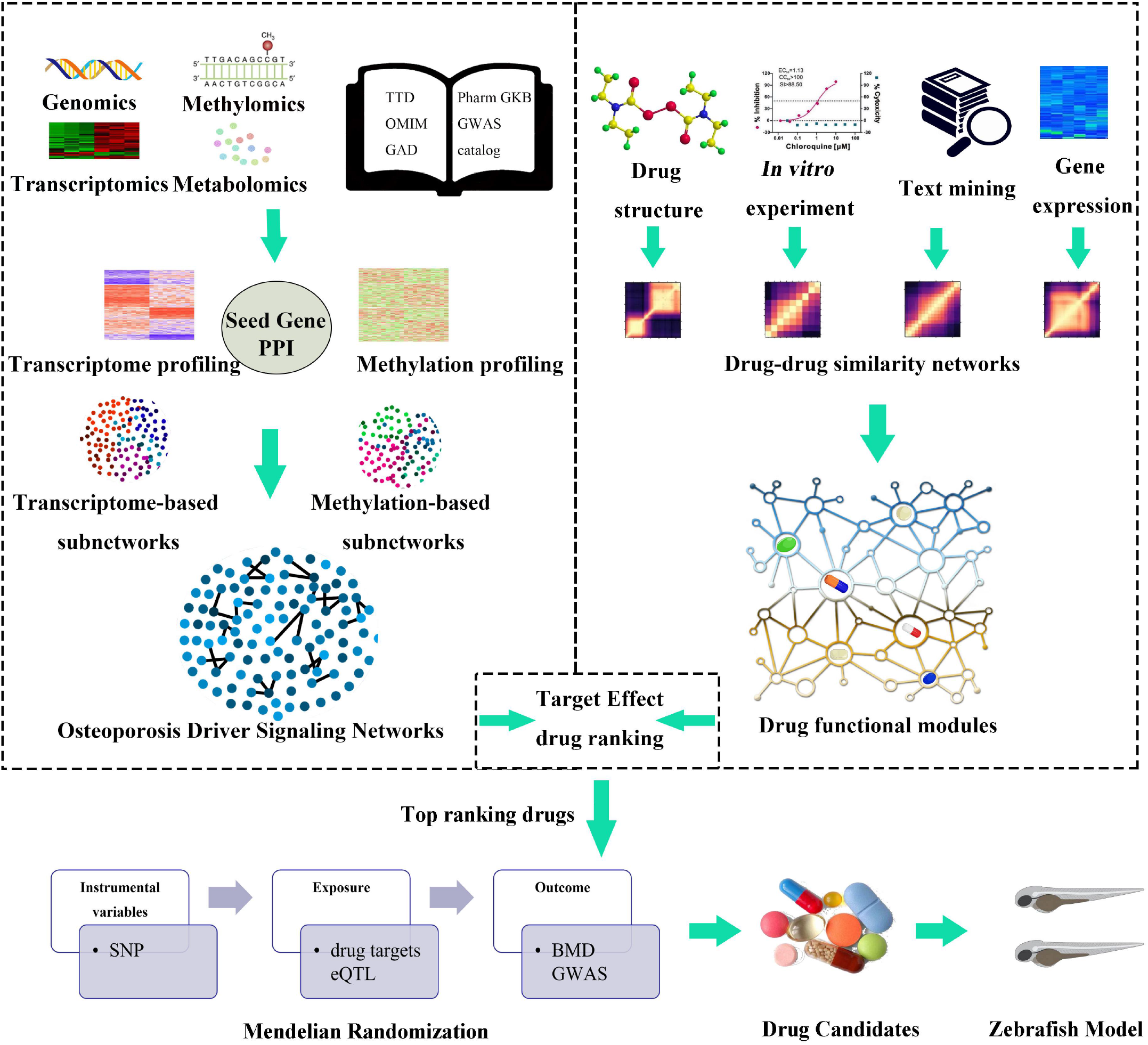
Overview of multi-omics based drug repositioning strategy. The systematic drug repositioning strategy included the following steps: (1) Osteoporosis driver signaling networks were identified from multi-omics data; (2) Drug functional modules were obtained using a network fusion approach to integrate drug similarity information; (3) All drugs were ranked according to targeting effects on driver signaling networks; (4) The causal relationships between drug targets and BMD were evaluated by MR analysis; (5) Potential drug repurposing candidates were validated in zebrafish model.

### 2.1 Construct osteoporosis-related driver signaling network

#### 2.1.1 Genes related to osteoporosis

Multi-omics data (dbGap phs001960.v1.p1) including genomics, methylomics, metabolomics, and transcriptomics were generated for 121 Caucasian female subjects using extreme phenotype sampling to select women with high (n = 62) and low (n = 59) BMD from the Louisiana Osteoporosis Study (Table 1) ^16–18^. The transcriptome and methylome profiles are from peripheral blood monocytes (PBMs), which are established to have functional genomic significance for bone metabolism ^19^. Our previous study has described the details of the inclusion and exclusion criteria, experimental procedures, multi-omics data sequencing, and quality control of omics data generation ^16^.

a. Transcriptomics: differential expression analysis of the RNA-seq data was performed using EdgeR ^20^. Differentially expressed genes (DEGs) were selected by comparing the gene expression data between high and low BMD groups at the threshold of *FDR*< 0.01 and |*logFC*| > 8 ^21^.
b. Methylomics: differentially methylated CpG sites (DMCs) were identified using logistic regression and the Sliding Linear Model (SLIM) method in methylKit ^22^. Significant DMCs with *FDR*< 0.05 were annotated to their corresponding genes.
c. Metabolomics: the two-sample t-test (*P-value* < 0.05) and partial least squares discriminant analysis (PLSDA) (Variable important in projection, *VIP-value* > 1.5) were performed to identify differentially abundant metabolites, which were then searched in HMDB (Human Metabolome Database) to determine the target genes associated with osteoporosis-related metabolites ^23^.
d. Genomics: the Knowledge-based mining system for Genome-wide Genetics studies (KGG) software was used to identify genes associated with BMD by gene-based GWAS analysis with *P* < 0.20 (Bonferroni Correction) ^24,25^.

To obtain more potential osteoporosis genes, we also searched public databases with “osteoporosis” as the keyword, including GWAS catalog (studies based on hip BMD of Caucasian adults) ^26^, TTD (Therapeutic Target Database) ^27^, OMIM (Online Mendelian Inheritance in Man) ^28^, GAD (Genetic Association Database) ^29^, and Pharm GKB (The Pharmacogenomics Knowledgebase) ^30^.

#### 2.1.2 Network-based Computational Strategy

Transcriptomic and methylation profiles were used to identify differentially expressed subnetworks with a higher network score compared with the background network based on osteoporosis-related genes by BMRF (Bagging Markov Random Field) tool ^31,32^. BMRF utilizes a simulated annealing method and follows a maximum a posteriori (MAP) principle to search subnetworks with maximal network scores ^31^. The background network was obtained from Pathway Commons (version 11), which contains data from several pathway databases and protein-protein interactome (PPI) databases with over 4,700 pathways and 2.3 million interactions ^33^. In the BMRF method, the significant score of one gene in a subnetwork depends not only on its expression profile or methylation profile from the multi-omics data, but also on the profiles of their neighbors in the PPI network ^32^. First, the subnetwork searching started from every osteoporosis-related gene (seed gene) that formed an initial subnetwork with a single node, then randomly sampled a new gene in the background network to get a new subnetwork. Genes adjacent to the current network can be added, and the subnetwork gradually increased a predefined network score *NetScore(G)*, which was defined as a negative posterior potential function measured from an estimated discriminative Z-score of the gene between low and high BMD groups ^32,34^. Simulated annealing stopped and outputted a final subnetwork when no significant improvement of the network score was achieved. Genes capable of forming a subnetwork with a more reliable frequency were designated as final likely driver genes based on these networks, for which the confidence score of each osteoporosis-related gene was computed by using a bagging scheme based on a bootstrapping strategy. The confidence score of gene *j* is calculated as follows:

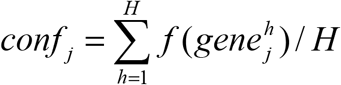

*h* is the number of bootstrap repetitions, *f*(*gene*) is 1 if gene *j* is selected in the *h*-th bootstrap replicate, and the program was run for 100 iterations. In the course of bootstrapping, the confidence level threshold was set to 0.2 to find significant genes in the network, as previously suggested ^15^.

After selecting the driver genes with high joint confidence scores, the methylation data-based signaling networks were merged with networks derived from transcriptomics data to construct a comprehensive driver signaling network. The AP (Affinity Propagation) algorithm, which is an unsupervised clustering approach that does not require *a priori* knowledge of cluster numbers, was applied to integrate the transcriptome/methylation-based osteoporosis subnetwork into the final osteoporosis signaling driver network based on the similarity between subnetworks ^35^. The similarity between transcriptome-based and methylation-based subnetworks was defined by the ratio of the shared genes between two subnetworks to total genes.

### 2.2 Drug functional modules based on drug-drug similarity network

Drug functional module construction aimed to identify drug communities that shared a common function in drug response and modes of action mechanisms ^36^. To construct drug functional modules, a non-linear network fusion technique (R package SNFtool v2.3.0) was used to integrate four independent drug similarity networks based on structure similarity ^37^, text mining similarity, *in vitro* test similarity, and drug-induced transcriptional response similarity, respectively ^38^. Two-dimensional chemical structure similarity (defined by the Tanimoto 2D chemical similarity scores), *in vitro* test similarity (defined by Pearson correlation of the activity patterns of the drugs based on NCI60 cell lines), and text mining similarity (based on co-occurrence scheme and natural language processing approach) between two drugs were extracted from the STITCH (Search Tool for Interacting Chemicals) database ^39,40^. Drug-induced transcriptional response data were extracted from the Library of Integrated Network-based Cellular Signatures (LINCS) database ^38^, which is a large consortium dedicated to creating a comprehensive reference library of cell-based perturbation-response signatures. To generate drug-induced transcriptional response similarity, a gene rank list was created for multiple doses of the drugs by the measure of the distance between two ranked lists based on their *logFC* value in response to drugs using Spearman’s Footrule and the Borda Merging Method ^41,42^. The Kruskal algorithm obtained a single rank list from a set of rank lists in a hierarchical way ^42^. Subsequently, each drug’s gene signature was determined by selecting the 250 genes that were ranked top and bottom for each drug according to their empirical distribution. The network-based gene set enrichment analysis (GSEA) score was used as the dissimilarity metric for the gene signatures between drug *i* and drug *j*. Finally, the AP algorithm was used to cluster the similarity network into drug functional modules.

Increasing evidence indicates that fat and muscle-derived myokines, adipokines, and growth factors could regulate skeletal remodeling and BMD, and thus in the pathogenesis of osteoporosis ^43^. Therefore, bone, muscle, and adipose cell lines derived from bone marrow (CD34), normal primary skeletal muscle cells (SKL) and normal primary adipocyte stem cells (ASC), respectively, in LINCS were selected because of the relationships among these tissues and fracture risk ^44–46^.

### 2.3 Drug repositioning based on osteoporosis driver signaling network (ODSN)

All candidates were ranked according to the evaluation of drug target effect on ODSN. The known drug-target interactions were extracted from public databases and the predicted drug-target interactions were determined based on the domain tuned-hybrid (DT-Hybrid) recommendation method ^47^. The strength of the targeting effects between each drug and ODSN were calculated by factorizing the drug-induced gene expression profiles into weight matrices and effect matrices using a Bayesian factor regression approach ^48,49^.

#### 2.3.1 Known drug-target and predicted drug-target

Known drug-target interactions were collected from SuperTarget, which is a comprehensive database of 332,828 drug-target interactions, and mapped by UniProt ^50–52^. Different information-theoretic concepts of functional similarity among genes have been developed for GO terms, including BP (Biological Process), MF (Molecular Function) and CC (Cellular Component) were calculated through the “GOSim” R package, and merged into a target-target similarity matrix by the AP algorithm ^53^. Then, the known drug-target interactions and drug functional network were used to predict off targets based on the DT-Hybrid recommendation algorithm ^47^. All known drug-target and predicted drug-target interactions were considered in our analysis.

#### 2.3.2 Drug-induced gene expression profile and target effects

A drug treatment transcriptional response matrix ***X*** (*n* × *m*) was derived from LINCS for each drug in every drug functional module, defined based on ***W*** (weight matrix) and ***λ*** (effect matrix). Each column of ***X***, i.e., ***X***_*i*_, is an *n* dimensional vector of gene fold-change (control vs. treatment) of drug *i* in the gene expression profile; *m* is the number of drugs in each drug module; *n* is the number of gene nodes in driver signaling network. By factorizing the treatment profiles, underlying signatures were generated using the BFRM model form ***X***_*i*_ = ***Aλ***_*i*_ + ***ε***_*i*_ (*i* = 1,2, ⋯, *m*) ^48,49^. Identify the networks involved in an unknown pharmacologic mechanism of a drug and to what extent they are related, a weight matrix was developed, ***W*** with *W_ij_* = *A_ij_* if *ρ_ij_* > *c* and *W_ij_* = 0 if *ρ_ij_* ≤ *c* (*c*=median *ρ*). c quantified the weight of gene *j* in the column of gene signature *k*, a matrix ***ρ*** quantified the probabilities of how each gene was associated with each factor ***λ***, and an effect matrix ***λ*** = (***λ***_1_, ***λ***_2_, ⋯, ***λ***_***m***_) with *λ_k,i_* quantified the effect of drug *i* imposed on the gene signature, ***E_k_***.

The quantified value of drug *d_i_* assessed to gene signature ***t*** is obtained by

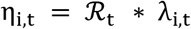

where 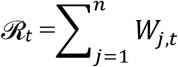 inferred by the response of gene signature ***t*** to the drug *_i_*.

For each weight matrix, a gene that was not a target of the drug *d_i_* was considered less meaningful and set equal to 0. Thus, for a driver signaling network *m_i_* and drug *d_i_*, ***η*_*midi*_** = (*η*_*g*1*midi*_, *η*_*g*_2*midi*, ⋯, *η*_*gkmidi*_), where {*g*_1_, *g*_2_ ⋯ *g_k_*} describes the elements of a network *m_i_*.

All the drugs were ranked according to the targeting effect score *TE*, which characterizes the effects of drugs on the derived driver signaling networks.

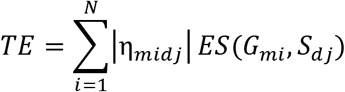

Here *N* is the number of osteoporosis driver signaling networks, |***η***_*midj*_| is the norm of ***η**_midj_* that indicates the targeting strength of drug *d_j_* on the driver signaling network *m_i_, G_mi_* is the gene set of disease on *m_i_*, *S_dj_* is the gene expression fold change of drug *d_j_*, and *ES*(*G_mi_, S_dj_*) corresponds to the enrichment score from GSEA ^54^.

### 2.4 Mendelian Randomization analysis of candidate drug target genes and BMD

Two sample Mendelian Randomization (TSMR) analysis was used to assess the potential causal association between the expression of drug targets and BMD using expression quantitative trait loci (eQTL) (sample 1) and published genome-wide association studies on BMD (sample 2) ^55^. For each drug candidate, the DrugBank database provided information about the target and actions (inhibitor or antagonist) with full descriptions, accessed on 10 May 2021 ^56^. We used drug target eQTL to mimic exposure to medications and downloaded the summary cis-eQTL results and allele frequency information derived from Genotype-Tissue Expression (GTEx) on 18 May 2021 ^57^. Significant cis-eQTLs (*P-value* < 5E-8) were selected as instrumental variables from specific tissues where the most significant (minimal *P-value*) eQTLs are observed. Variants related to bone mineral density were sourced from the IEU OpenGWAS project, which contained >200 billion genetic associations from >40,000 GWAS summary datasets ^58^. By searching “bone mineral density” as a keyword in the trait of the IEU OpenGWAS project, 13 GWAS summary datasets generated by different consortia were collected as outcomes for further MR analysis (Supplemental Table S2), and detailed recruitment and quality control are available in the previous publications. The inverse variance weighted (IVW) method was used to estimate the causal effect between the exposure (i.e., expression of drug target) and outcome (i.e., BMD), as previous studies have shown that IVW has the highest test efficiency among the various MR estimation methods ^59,60^. The significance threshold was determined using the Bonferroni method to adjust for multiple hypothesis testing. Sensitivity analyses were conducted to test for heterogeneity in the causal effects estimated with the different instrumental variables, and to rule out the presence of horizontal pleiotropic effects.

### 2.5. Preventive Effects of candidate drugs on Dexamethasone-Induced Osteoporosis in Zebrafish

To further evaluate the effect of predicted candidate drugs on bone mineralization, we established dexamethasone (Dex)-induced zebrafish osteoporosis model and exposed zebrafish larvae to different concentrations of each drug candidate. Dex-induced zebrafish is a commonly used glucocorticoid-induced osteoporosis model for evaluating drug efficacy in drug discovery, in which drug treatments that alleviate Dex-induced osteoporosis in zebrafish are thought to activate bone remodeling and be effective for the treatment of osteoporosis ^61,62^.

#### 2.5.1 Reagent

Dex and Alfacalcidol (AC) were purchased from Sigma-Aldrich (Darmstadt, Germany). Amiloride HCl, Aminocaproic acid, Acenocoumarol, and Acebutolol HCl were purchased from Abmole Bioscience (Houston, USA).

#### 2.5.2 Animals

The wild-type TU (Tuebingen) zebrafish were provided by the Department of Genetics and Development Biology, College of Life Sciences in Hunan Normal University, and bred in natural pairs. The Hunan Normal University Institutional Animal Care and Use Committee approved the animal experiment protocols. At 28.5 °C and 14/10 h light/dark cycles, Zebrafish embryos and adults were cultured in an aquatic recirculating system and were fed twice per day according to the standard guide for the laboratory use of zebrafish ^63^.

#### 2.5.3 Experimental procedures

Zebrafish embryos were randomly divided into 7 groups (n = 20 embryos/2 wells/group) as follows: 0.1% DMSO (Dimethyl sulfoxide, control group), 10 μmol/L Dex (Dexamethasone, model group), 10 μmol/L Dex+ AC (Alfacalcido, positive control group, 10, 1, 0.1, 0.01 μg/ml), 10 μmol/L Dex+ Amiloride HCl (10, 1, 0.1, 0.01 μg/ml), 10 μmol/L Dex+ Acebutolol HCl (10, 1, 0.1, 0.01 μg/ml), Aminocaproic acid (10, 1, 0.1, 0.01 μg/ml), Acenocoumarol (10, 1, 0.1, 0.01 μg/ml) from 3 dpf (day post fertilization) to 9 dpf. The culture water was changed in each 12 hours at 0 dpf to 3 dpf (Table 2). The half Drug-containing culture water were changed in each 24 hours at 3dpf to 9 dpf. All groups were dissolved into a 0.1% DMSO mixture.

#### 2.5.4 Alizarin red staining

Zebrafish larvae were collected and fixed in 2% paraformaldehyde for one hour at 9 dpf. After washing with 100mM Tris /10mM MgCl_2_, bleach was performed with 3% H_2_O_2_/0.5% KOH. Next, these fish larvae were rinsed with 25% glycerol/0.1% KOH until no bubble generation. After removing the rinsing liquid, the larvae were stained with 0.01% Alizarin red stain for one hour, with 50% glycerol/0.1% KOH to stain formed bone for 10 minutes and subsequently destained with 50% glycerol/0.1% KOH. Finally, digital photographs were taken using a stereomicroscope.

#### 2.5.5 Quantitative analysis of mineralized bone

The zebrafish of the ventral side stained with Alizarin Red was selected for observation under a stereomicroscope, and images were acquired with imaging software (100×). All images were acquired under the same light intensity and exposure settings. Image J was used to select the alizarin red-stained area by setting a threshold, and its area was calculated to reflect the amount of bone mineralization.

#### 2.5.6 Statistical analysis

One-way analysis of variance (one-way ANOVA) was used for the comparison between groups of normally distributed data, with *P* < 0.05 indicating statistical significance in multiple groups. The LSD (Least Significant Difference) method was used for multiple comparisons between groups when the variances were homogeneous, otherwise, the Dunnett T3 method was performed ^64^.

## 3. Results

### 3.1 Osteoporosis Driver Signaling Network (ODSN) Identification

To identify the ODSN, we first obtained an osteoporosis-related gene set based on multi-omics data. We identified 359 mRNAs as differentially expressed genes (*FDR*< 0.01, |logFC| > 8.00), 169 genes with 397 differentially methylated sites (FDR < 0.05), and 19 mapped genes from differentially abundant metabolites. Two genes (BCL2 and BASP1) were prioritized by gene-based GWAS analysis at the threshold of *P* < 0.20. Furthermore, 14 genes from TTD, 295 genes from GAD, 88 genes from the GWAS catalogue, 39 genes from OMIM, and 9 genes from PK were identified as osteoporosis-related genes. After removing duplicates, there were 944 potential driver genes to be used as seed genes for BMRF (Supplemental Table 3). After removing genes not included in the PPI network, 586 and 605 seed genes were selected for constructing transcriptome- and methylation-based networks, respectively.

559 mRNA-based subnetworks and 538 methylation-based subnetworks were identified by using the BMRF algorithm (Supplemental Table 4). After merging the methylation-based subnetworks with transcriptome-based subnetworks by selecting the genes with high joint confidence scores (threshold = 0.20) and using the AP algorithm to cluster the osteoporosis driver networks, 206 subnetworks were constructed and used for further analysis (Supplemental Table 5).

### 3.2 Drug Functional Modules Construction and Target Effect

169 drugs and their similarity scores formed a similarity network by using the nonlinear network fusion technique to integrate four drug similarity networks into one uniform network. After clustering by the AP algorithm, 169 drugs were grouped into 33 drug functional modules through similarity scores in each cell line (Supplemental Table 6). SuperTarget identified 2469 interactions containing 72 drugs and 1367 targets. Finally, 16864 interactions containing 1367 targets of 113 drugs were predicted by DT-Hybrid recommendation algorithm (Supplemental Table 7).

### 3.3 Drug Ranking

The potential of all drugs to treat osteoporosis was predicted by evaluating the targeting effect of each drug on the ODSN in each cell line, and we ranked all the drugs according to the *TE* correlation score (Table 3). Finally, we identified 8 drugs as potential therapeutic candidates since they ranked in the top 20 drugs in bone/muscle/adipose cell lines. These candidates included Amoxapine (anti-depressant), Armodafinil (wake-promoting agent), Acebutolol (anti-hypertensive), Acenocoumarol (anti-coagulant), Amiloride (diuretic), Abacavir (anti-retroviral), Artesunate (anti-malarial) and Aminocaproic acid (anti-fibrinolytic).

### 3.4 MR analysis to estimate the association of drug targets with BMD

To estimate the potential effect of drug candidates on osteoporosis, we used a TSMR analysis to determine the association between the expression of drug targets and BMD. Five drug target gene expressions were detected to be associated with BMD, including Heel BMD, Femoral neck BMD, Lumbar spine BMD, and Forearm BMD (Figure 2, Table 4, Supplemental Table 8).

**Figure 2.**
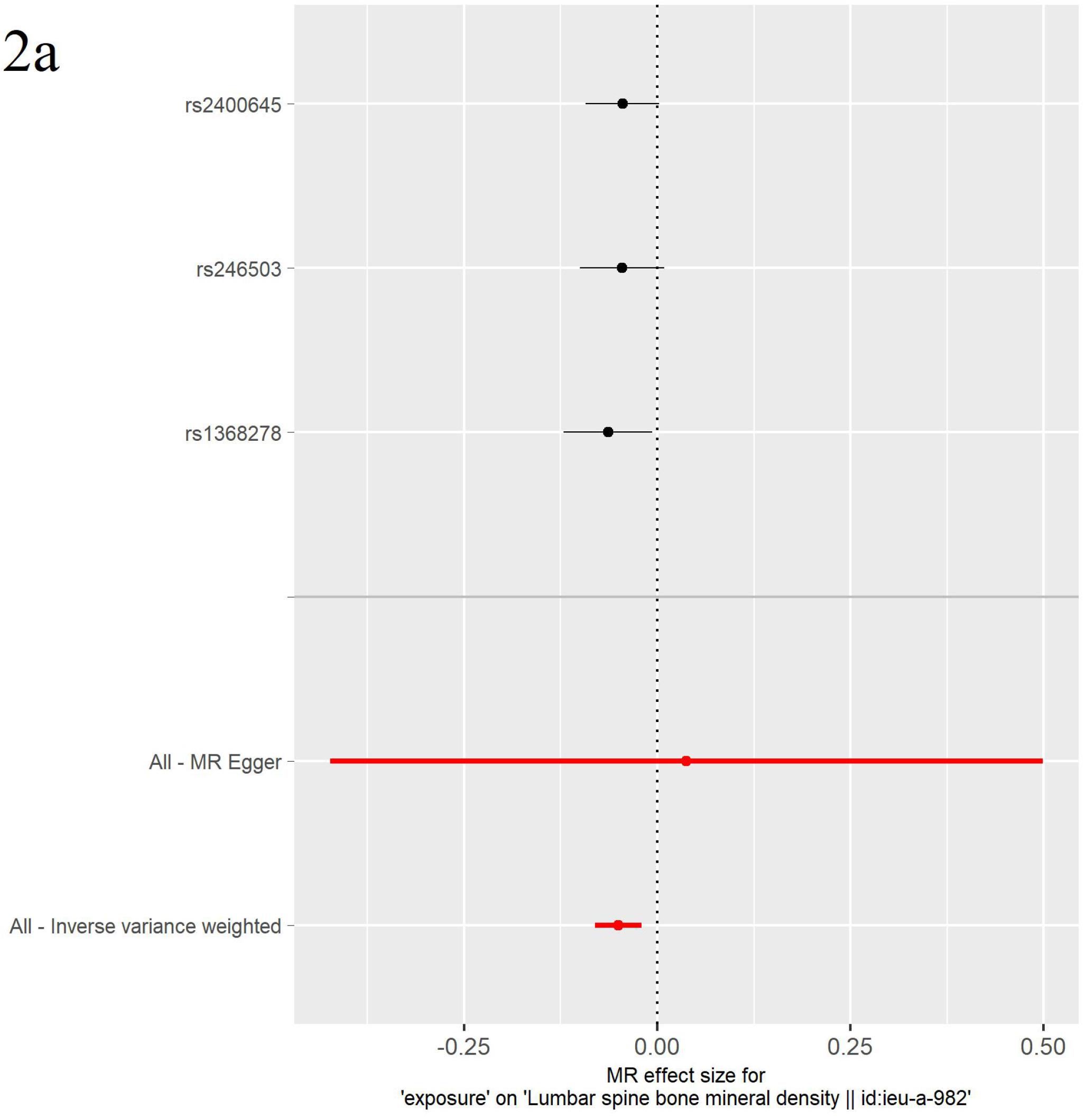
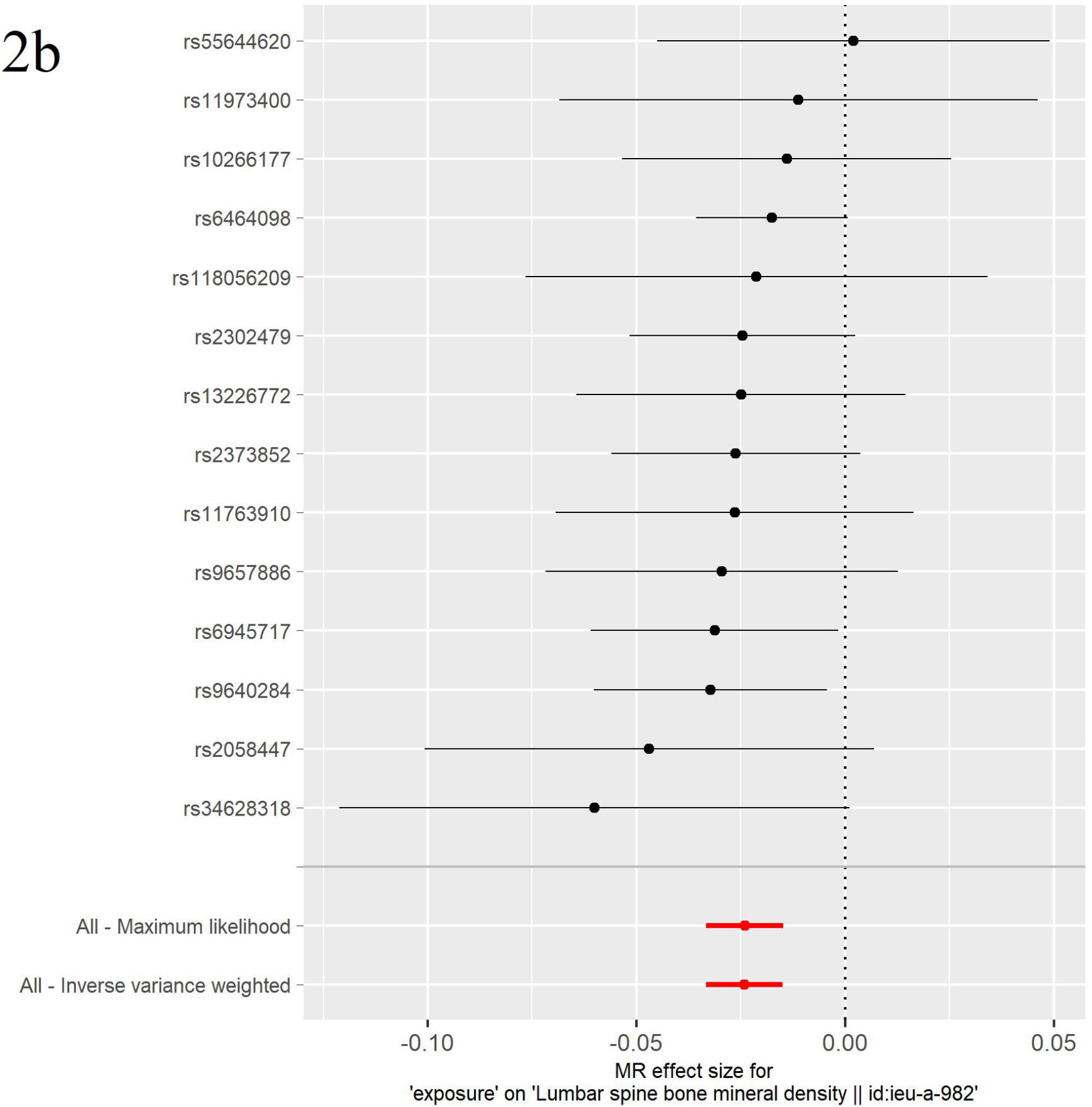
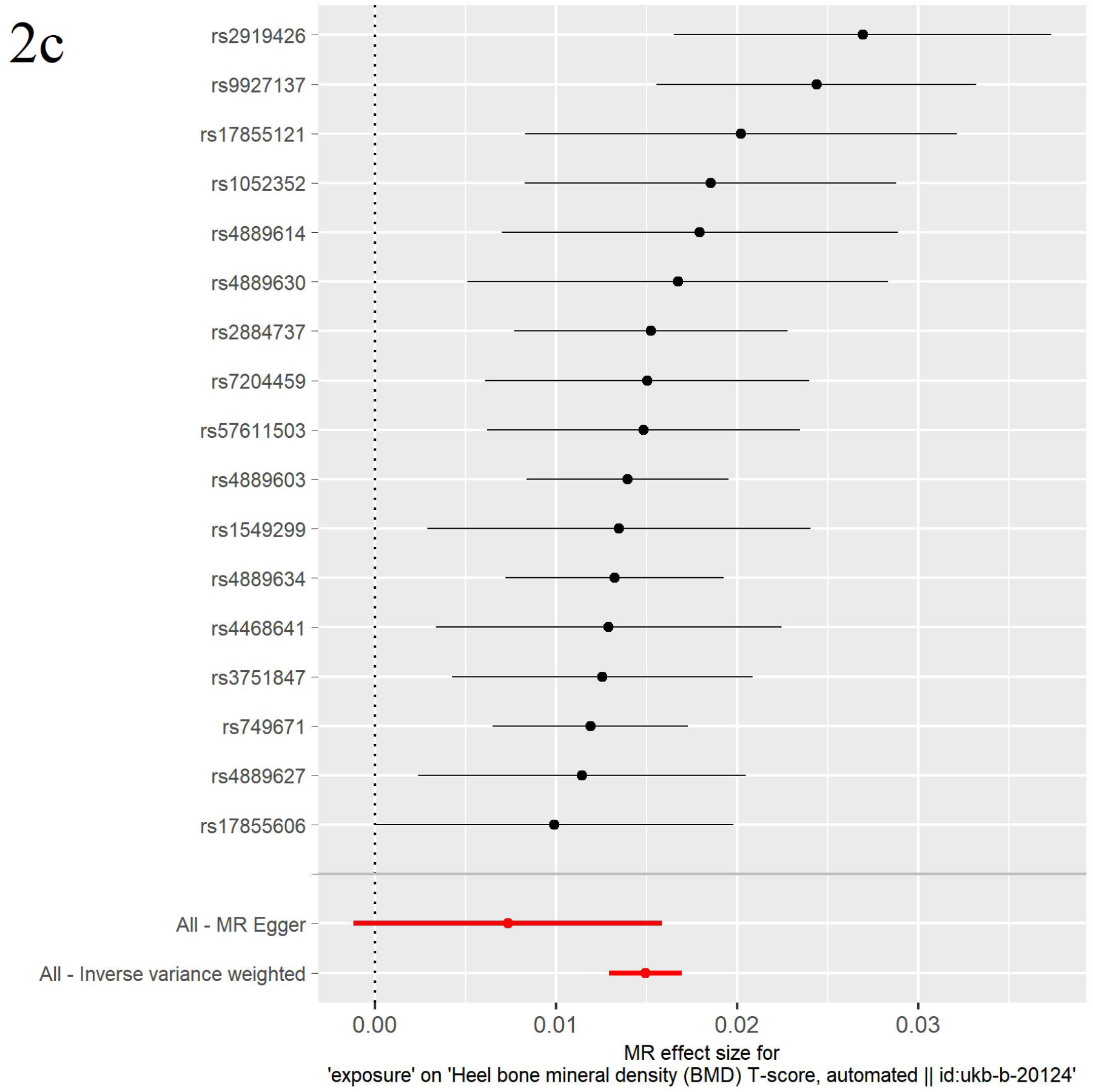
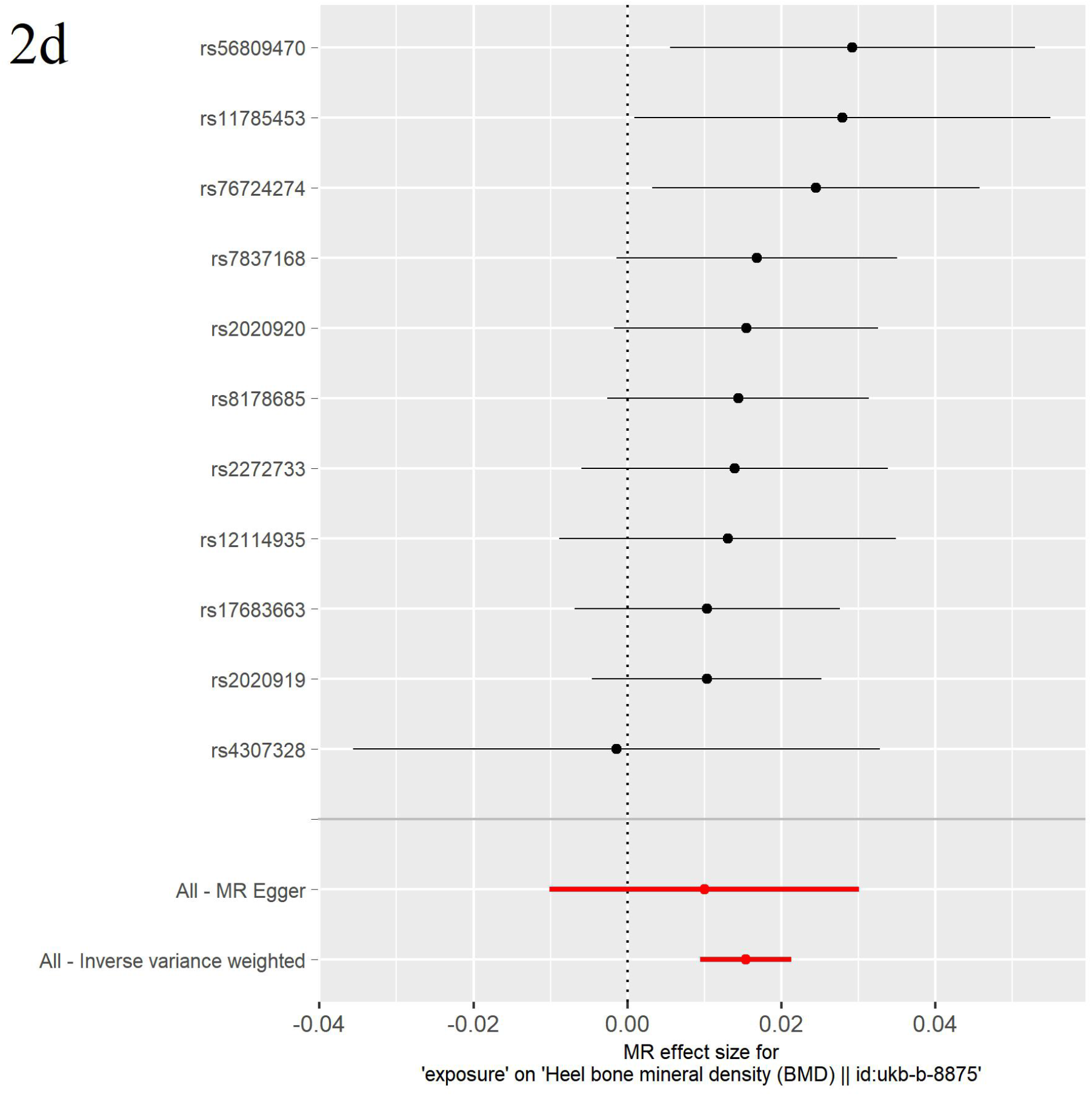
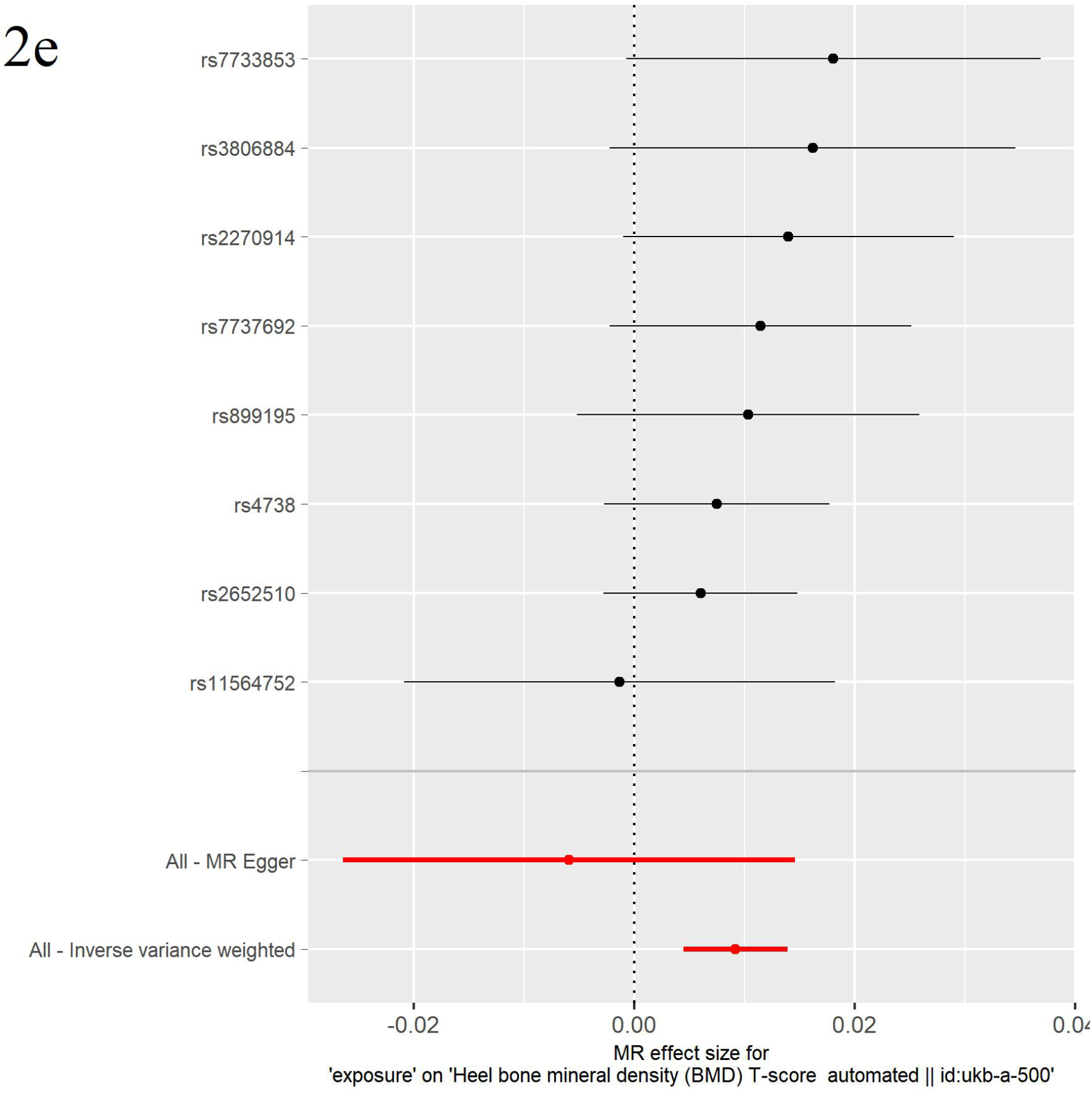
Forest plots showing the effects of alleles associated with drug target genes expression on the risk of BMD. Mendelian Randomization analysis showed that (2a) Acebutolol and (2b) Amiloride through inhibiting expression of target ADRB2 and AOC1, respectively, were significantly associated with increased BMD. (2c) Acenocoumarol, (2d) Aminocaproic acid and (2e) Armodafinil were detected to decrease BMD.

#### Acebutolol and Amiloride

Acebutolol and Amiloride were hypothesized to be associated with increased BMD based on the effects of their known targets. For ADRB2 (the target of Acebutolol), a total of 3 cis-eQTLs in atrial appendage tissue were used in the TSMR analysis. IVW results showed the estimated effect of ADRB2 expression, which is inhibited by Acebutolol, on BMD was negative (*β* = −0.05, *P*-value = 1.06E-3, Figure 2a). We observed that there was no significant heterogeneity among the instruments (*P*-value = 0.29) and horizontal pleiotropic effects (*P*-value = 0.77). The leave-one-out test showed that removing any SNP would not have a fundamental impact on the outcome, suggesting that the MR results were robust. Amiloride inhibits expression of AOC1, which was also observed to have a negative effect on BMD using cis-eQTLs from whole blood tissue (*β* = −0.02, *P*-value = 2.03E-7, heterogeneity *P*-value = 0.98, pleiotropy *P*-value = 0.34, Figure 2b).

#### Acenocoumarol, Aminocaproic acid and Armodafinil

Acenocoumarol, Aminocaproic acid and Armodafinil were hypothesized to promote bone loss. Acenocoumarol inhibits VKORC1, which was observed to have a positive effect on BMD (*β* = 0.01, *P*-value = 2.92E-48, heterogeneity *P*-value = 0.53, pleiotropy *P*-value = 0.09, Figure 2c) using cis-eQTLs from liver tissue. Aminocaproic acid inhibits expression of PLAT, which had a positive effect on bone (*β* = 0.02, *P*-value = 3.38E-7, heterogeneity *P*-value = 0.91, pleiotropy *P*-value = 0.59, Figure 2d) using cis-eQTLs from skin tissue. Armodafinil inhibits expression of SLC6A3, which was found to have a positive effect on BMD (*β* = 0.01, *P*-value = 1.40E-4, heterogeneity *P*-value = 0.82, pleiotropy *P*-value = 0.19, Figure 2e) using cis-eQTLs from lung tissue.

## 3.5 Effect of candidate drugs on bone mineralization in Dex-induced osteoporosis zebrafish model

A Dex-induced osteoporosis zebrafish model was implemented to further evaluate the predicted beneficial or harmful effects of the candidate drugs. Dex treatments (10 μmol/L) reduced bone mineralization of Dex-induced zebrafish in contrast with normal zebrafish samples, indicating that the osteoporosis model was successfully established (Figure 3a-3b, Figure 4, Table 5). Amiloride HCl at doses of 10 μg/ml and 1 μg/ml significantly increased bone mineralization in Dex-induced osteoporosis zebrafish (Figure 3c-3d). Acebutolol HCl at doses of 10/1/0.1/0.01 μg/ml significantly increased bone mineralization in Dex-induced osteoporosis zebrafish, and doses of 10/1/0.1 μg/ml improved a larger bone area compared with 0.01 μg/ml (Figure 3e-3h). Compared with the positive control group (10/1 μg/ml Alfacalcido treatment zebrafish died during 3-9dpf), Acebutolol HCl with 0.1/0.01 μg/ml had no significant difference in bone mineralization compared with Alfacalcido at doses of 0.1/0.01 μg/ml (Figure 3i-3j). Aminocaproic-acid (10, 1, 0.1, 0.01 μg/ml) treatment did not show significantly enhanced or reduced bone mineralization compared with the blank control group (0.1% DMSO). Acenocoumarol (0.1 μg/ml) reduced the mineralization area compared with the blank control group (Figure 3k).

**Figure 3.**
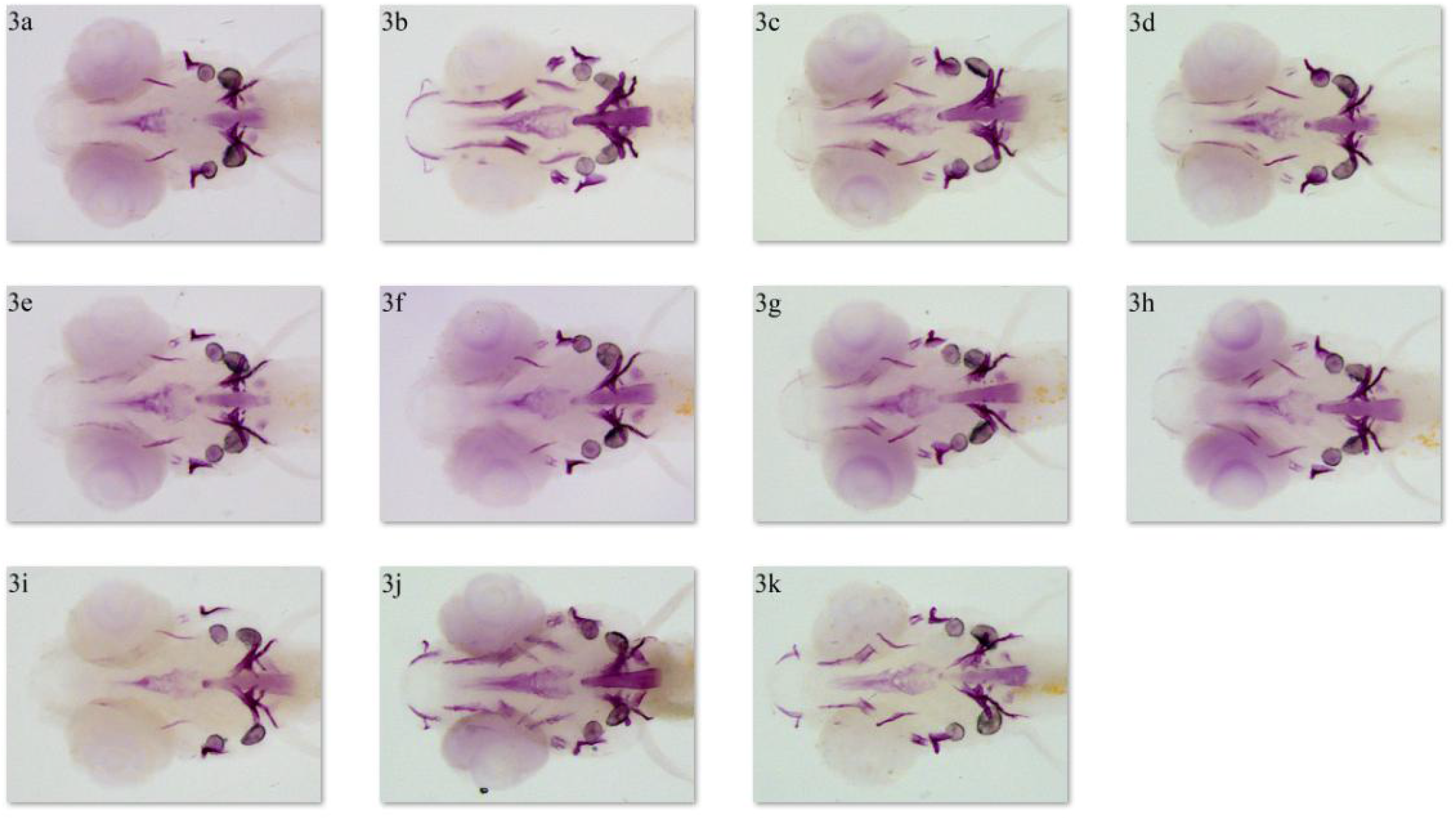
Alizarin red staining for different concentrations of drug treatment in zebrafish larvae at 9 dpf. The results showed the alizarin red staining area of zebrafish skulls in the model group (3a), the control group (3b), the Amiloride HCl groups at a dose of 10 μg/ml (3c) and 1 μg/ml (3d), Acebutolol HCl groups at the dose of 10/1/0.1/0.01 μg/ml (3e/3f/3g/3h), Alfacalcido groups at the dose of 0.1/0.01 μg/ml (3i/3j) and Acenocoumarol group at the dose of 0.1μg/ml (3k).

**Figure 4.**
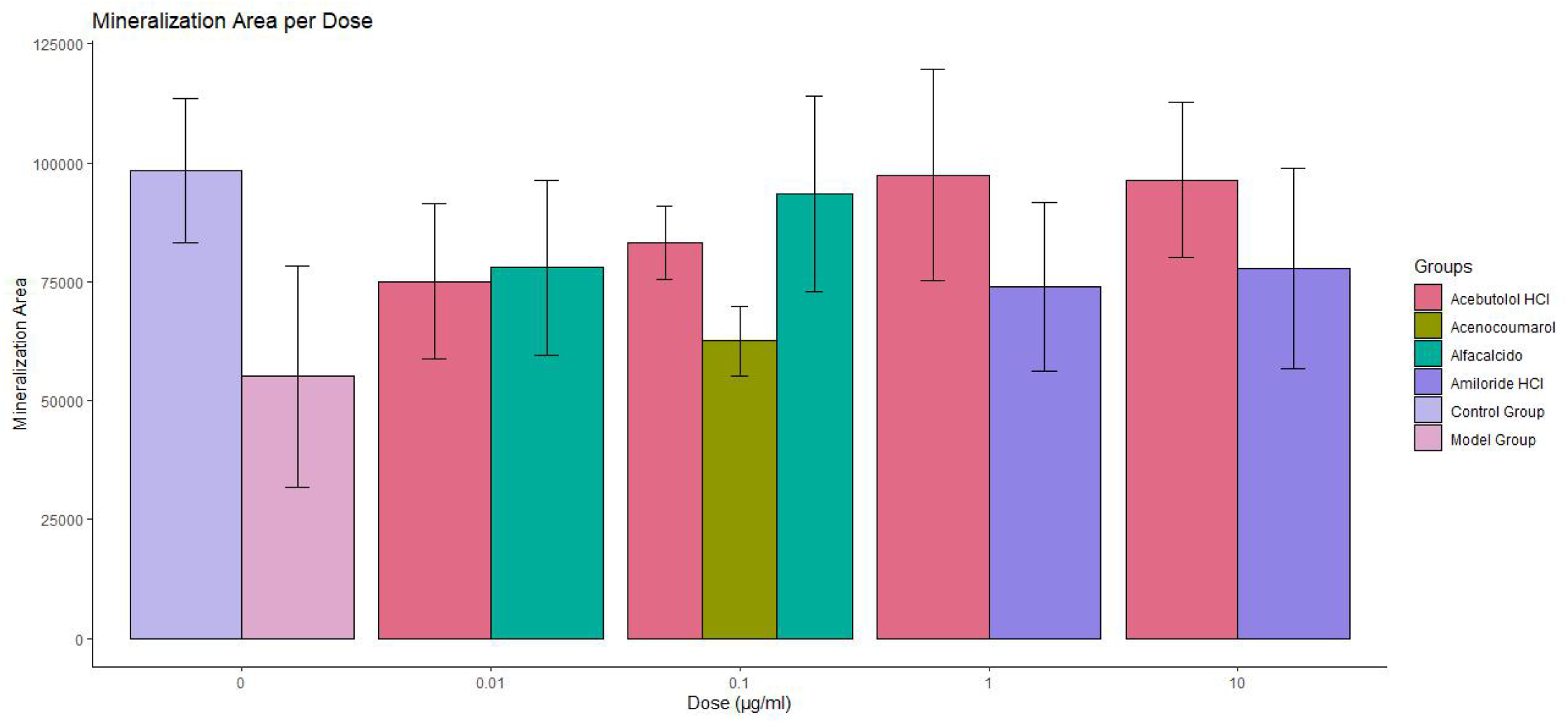
Mineralization area of drug treatment per dose. All values are expressed as means ± SDs.

## 4. Discussion

This drug repurposing study aimed to assess the innovative adaptation of existing drugs for novel osteoporosis treatment. ODSN and DFN were established, and target effect scores were computed to select the potential drugs for osteoporosis. MR was then used to predict the positive or negative drug effect on disease and to suggest the potential target mechanism. Five drug candidates were identified and recommended based on their high-rank scores and significant causal relationships of their targets with BMD variation. Acebutolol HCl and Amiloride HCl were observed to improve bone mineralization area in the Dex-induced osteoporosis zebrafish model, and Acenocoumarol reduced the bone mineralization area. Thus, Acebutolol and Amiloride might enable the development of potentially novel treatment agents for osteoporosis.

Acebutolol, a cardio-selective beta-adrenergic blocking agent, and Amiloride, a diuretic, are both commonly used for the treatment of hypertension. In hypertension patients, there is an increase in urinary calcium excretion, leading to an increase in the release of calcium from the bones, which may in turn accelerate osteoporosis ^65^. Numerous studies have also provided evidence that BMD and cardiovascular disease are influenced by common factors such as bone morphogenetic proteins, PTH, and dyslipidemia ^66^. Additionally, studies have demonstrated that the processes of osteogenesis and angiogenesis are interconnected during bone remodeling ^67^. The primary mode of action for Acebutolol is to inhibit ADRB1 (β1-adrenergic) and ADRB2 (β2-adrenergic) receptors. Although Acebutolol has not previously been recommended for osteoporosis treatment, the effect of beta-blockers on fracture risk has been widely discussed ^68,69^. Investigators have reported that adrenergic agonists may influence bone metabolism by stimulating bone resorption ^69^, increasing osteoclastogenesis *in vitro*, and regulating osteoclast activating cytokines ^70^. ADRB1 signaling was reported to regulate bone anabolic responses during growth and in response to loading, and ADRB2 was reported to inhibit bone formation and promote osteoclastogenesis by increasing RANKL ^71,72^. While the underlying mechanisms of the relationship between beta-blockers and bone metabolism are still not well understood ^68^, a beta-1 adrenergic blocker Atenolol is currently being investigated in Phase 3 randomized clinical trial for the prevention of bone loss (ClinicalTrials.gov ID: NCT04905277). On the other hand, Amiloride is a diuretic used in combination with other therapeutic agents to treat hypertension on a long-term basis ^73^, which was previously reported to inhibit osteoclastogenesis by suppressing nuclear factor-kB and mitogen-activated protein kinase activity ^74^, supporting the reliability of our findings.

Acenocoumarol (anti-depressant agent), Aminocaproic acid (anti-fibrinolytic agent), and Armodafinil (wake-promoting agent) may potentially reduce BMD, implying that the long-term effects of these medications on osteoporosis risk need vigilant monitoring. It was reported that the mean decrease in femur BMD was 1.8% and 2.6% in 86 patients treated with Acenocoumarol for 1 and 2 years of follow-up, respectively ^75^. Acenocoumarol may potentially promote osteoporosis by inhibiting VKORC1 (Vitamin K epoxide reductase complex subunit 1), which is involved in vitamin K metabolism and normal bone development ^76^. Young male mice treated with modafinil (enantiomer of Armodafinil) developed trabecular and cortical bone loss, and their biomechanical strength was decreased ^77^. Although the possible mechanism of Aminocaproic acid in osteoporosis has not been reported, Aminocaproic acid is an antifibrinolytic drug may affect BMD through its effect on PLAT (Tissue-type plasminogen activator) ^78,79^, as osteocalcin was recently reported to be released from hydroxyapatite by plasmin ^80^.

Despite the novelty of this drug repurposing analysis in the bone field, several limitations should be taken into consideration. First, the multi-omics samples are derived from 121 normal Caucasian women, which, though one of the largest samples sizes available so far in the field, is still moderate and may render less comprehensive detection of the osteoporosis driver networks compared with larger sample sizes. Second, the LINCS project examined the expression of just 978 genes and imputed the expression of the remainder of the genome. Therefore, these changes in gene expression may not be indicative of the real mechanisms of action of some compounds, which might cause a certain unknown degree of bias. Third, we excluded non-overlapping multi-target effect drugs in the MR study because potential multi-target and pleiotropic effects of compounds cannot be ruled out and quantified entirely. Since osteoporosis is a complex multifactorial disease, a single target effect on BMD is lacking comprehensive mechanisms to be illustrated. How multiple targets affect the development of osteoporosis is a critical question, and more robust evidence needs to be considered on multi-target drug mechanisms for osteoporosis treatment. Furthermore, Acebutolol and Amiloride were administered in the hydrochloride hydrate form in the zebrafish validation experiments due to solubility, which may result in some biopharmaceutic and pharmacokinetic heterogeneity compared to Acebutolol and Amiloride. To verify the accuracy of our prediction and elucidate the anti-osteoporosis mechanism from a perspective of pharmacology, future biological validation experiments that investigate how drugs affect normal bone cells such as osteoblasts and osteoclasts will be critical.

In summary, the present study suggested potential drug candidates for osteoporosis treatment and proposed bone loss pharmacovigilance with long-term use of others. Acebutolol and Amiloride were recommended as possible novel therapeutic options for osteoporosis. Acenocoumarol, Armodafinil, and Aminocaproic acid may decrease BMD and increase the risk of osteoporosis. Zebrafish experiment results offered meaningful evidence on drug treatment in osteoporosis. We hope that the findings from this study will help guide and accelerate research in osteoporosis drug development. However, the identified drug candidates still require future experimental validation and large-scale clinical trials before their use in osteoporosis management.

## 5. Acknowledgment

This study was supported in part by the Natural Science Foundation of China (NSFC; 81570807, 31970504 and 31772548). HWD was partially supported by grants from the National Institutes of Health [U19AG05537301, R01AR069055].

## 6. Author contributions

Conceptualization, Hong-Wen Deng, Li-Jun Tan, Yun Deng and Hong-Mei Xiao; methodology, Dan-Yang Liu, Qiao-Rong Yi, Shuang Liang, Yue Zhang, Jia-Chen Liu and Xiang-He Meng; software, Dan-Yang Liu; writing—original draft preparation, Dan-Yang Liu; writing—review and editing, Dan-Yang Liu, Jonathan Greenbaum, Li-Jun Tan and Hong-Wen Deng; visualization, Dan-Yang Liu; supervision, Li-Jun Tan and Hong-Wen Deng; funding acquisition, Yun Deng, Li-Jun Tan and Hong-Wen Deng. All authors have read and agreed to the published version of the manuscript.

